# GenotypeTensors: Efficient Neural Network Genotype Callers

**DOI:** 10.1101/338780

**Authors:** Joshua Cohen, Manuele Simi, Fabien Campagne

## Abstract

We studied the problem of calling genotypes using neural networks. A machine learning approach to calling genotypes requires a training set, an approach to convert genomic sites into tensors and robust model development and evaluation protocols. We discuss each of these components of our approach and compare four types of neural network training protocols, two fully supervised and two semi-supervised approaches. Semi-supervised approaches use unlabeled data to supplement limited quantities of labeled data. Random hyper-parameter searches identified highly performing models that reach indel F1 of 99.4% on a chromosomes 20, 21, 22 and X of NA12878/HG001. We further validate these models by evaluating performance on HG002, an independent sample used in the PrecisionFDA challenge. We apply GenotypeTensors to evaluate the impact of (1) training with small datasets, (2) training models only with sites inside confidence regions, or (3) training with improved true label annotations. A PyTorch open-source implementation of GenotypeTensors is available at https://github.com/CampagneLaboratory/GenotypeTensors. DNANexus cloud applications are provided to help process new datasets both to train model or call genotypes with trained models.

## INTRODUCTION

DNA can be sequenced with high-throughput sequencers to yield short or long reads. Genotype callers are tools that implement computational approaches to transform sequencing reads aligned to a genome into genotype calls across the sites of a genome. Several approaches have been followed to develop genotype callers. Early stage ad-hoc callers used hard thresholds and filters to identify the consensus of bases seen at each site. These approaches were quickly superseded by approaches that leverage probabilistic methods to weight the evidence for each genotype at a site. Examples of probabilistic callers include the Genomic Analysis Toolkit (GATK) [McKenna et al., 2010], or Platypus [Rimmer et al., 2014].

At the heart of the probabilistic approaches are models that given observables at each site of the genome determine the probability of each genotype at the site. The probabilistic tests implemented in traditional callers use a limited amount of observable features. For instance, they may rely on the number of bases supporting a given genotype and the quality scores of these bases. The reason for the focus on a small number of features is that it is non-trivial to manually develop models that model interactions, often non-linear, among many kinds of observables.

Poplin et al. [2016] and our group [Torracinta and Campagne, 2016] have independently developed approaches to estimate the probability function for genotype calling using features observed from alignment data and neural network probability estimators. The approach used to develop DeepVariant [Poplin et al., 2016] consists in transforming aligned reads into images and train convolutional neural networks to predict genotypes. In contrast, our approach maps aligned reads to a vector of features and uses feed forward network architectures. We describe this approach in this manuscript and provide details of a robust model development and evaluation process that allows to train strong models at a fraction of the computational cost of the approach implemented in DeepVariant.

A key challenge faced when training genotype callers is access to large amounts of high-quality labeled data, that is, alignments where the genotypes at each site of the genome are known. In this work, we used data measured from NA12878/HG001, one of the samples studied by the platinum genome study [Eberle et al., 2016], subsequently studied by the Genome in a Bottle (GIAB) Consortium [Zook et al., 2014, 2018]. In order to test if models trained on one genome generalize to a second genome, we use sample HG002, which was used as benchmark sample in the PrecisionFDA challenge. This evaluation protocol makes it possible to compare the performance of our approach to the performance reported for other genotype callers in the same condition.

Importantly, GIAB defined confidence regions as a subset of the genome where confidence in genotype calls is strong [Zook et al., 2014, 2018]. Genotypes for sites outside of a confidence region may be correct, but there is less evidence that they are correct, either because data are lacking at the site, or because complexity of the genomic site yields conflicting genotypes when different sequencing technology are used to measure the same site, or across pedigrees. We use confidence regions for evaluation on both HG001 and HG002. The DeepVariant preprint reported training models with sites only contained in confidence regions. In our initial preprint, we warned that this could yield biased models that do well inside confidence regions (where the evaluation is performed), but fare worse in the rest of the genome (on the more complex sites that make up about 14% of the genome).

In this manuscript, we test whether the performance of models trained exclusively on sites located inside confidence regions is representative of performance of models trained with sites from across the entire genome.

Since high-quality genotype calls are rare and millions of these calls are necessary to train strong models, we evaluate semi-supervised training protocols to determine if using large amounts of unlabeled data can be beneficial when training genotype callers. This question is particularly interesting because the performance of neural networks has been shown to increase with larger training sets, but the cost of acquiring millions of high-quality genotype labels is expected to remain very large in the foreseeable future. Semi-supervised approaches could facilitate training strong models with limited amounts of labels. Our results show that semi-supervised approaches do not result in more predictive models when testing on an independent sample.

Domain adaptation techniques have been developed to minimize performance drops when a model is trained on a dataset with different distribution as the test set. We therefore also evaluated a domain-adaptation approach: Adversarial Discriminative Domain Adaptation (ADDA) [Tzeng et al., 2017]. We found that ADDA did not improve HG002 performance when models trained on HG001 where adapted to the HG002 data distribution.

Further analysis revealed that the performance of models trained on HG001 generalize well to the validation set and test set of both HG001 and HG002, suggesting that differences in performance across the two genomic samples cannot be explained by overfitting.

Recent work showed that careful tuning of baseline architectures can yield state of the art performance compared to more complex architectures [Merity et al., 2018] (authors studied sequence models for natural language processing tasks). This study confirms that hyper-parameter tuning is critical to training state of the art neural network models. In practice, selecting optimal hyper-parameters is difficult because of the computational burden of training many models with different hyper-parameters. In this study, we present and take advantage of an approach that greatly speeds up hyper-parameter searches when the models are small and many models can fit in the memory of a single graphical processing unit (GPU).

## RESULTS

### Data Preparation

Figure 1 presents an overview of the process we followed to prepare data for neural network training. Briefly, short reads were aligned to the human genome, alignments were processed with HaplotypeCaller [McKenna et al., 2010] to realign SNPs in the proximity of indels and to reduce the dimensionality of the dataset to regions likely to contain variation. Alignments were converted to a vectorial representation suitable to train a feed-forward neural network. Figure 1 also illustrates the funnel architecture, which allows for interactions of every feature with every other feature and progressively reduces the size of the representation by applying a reduction factor r (see methods). The advantage of the funnel architecture is that it does not require a preconceived notion about which feature interactions should be considered. Training determines which combination of feature values is informative to predict the labels by adjusting the weights of the neural network. The feature mapper used in this work considers 5 possible genotypes for every position of the genome, and produces 2,255 features per genomic site (451 features per genotype at each genomic site). The features are very sparse (most values are zero), since for instance, a diploid genome is expected to have one or two genotypes present at each site (when a genotype is not observed in a sample, all 452 features of this genotype have value zero). Throughout this study, we observed that networks with 2 layers are sufficient to yield top performing models (see parameters of best performing models shown in Methods).

**Figure 1.**
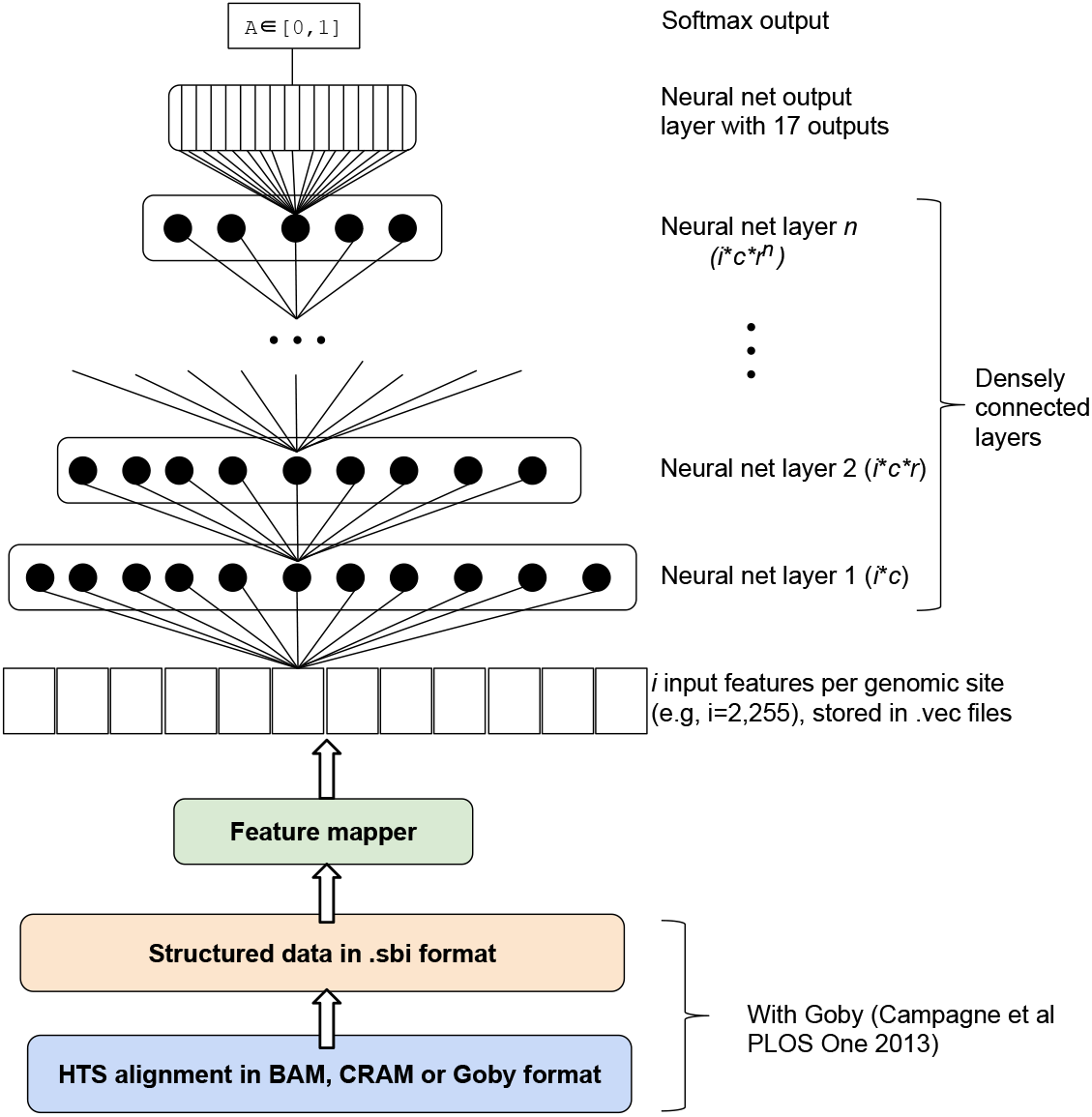
Overview of data generation, data management and neural network architecture. The process of developing a genotype caller model starts with HTS reads. The reads are aligned to the genome (here with BWA-mem, but other aligners can be used for different read characteristics). The BAM alignment is realigned with the GATK3 HaplotypeCaller [McKenna et al., 2010]. Realigned reads are converted to Goby format and converted to the Sequence Base Information format (extension .sbi). A feature mapper converts data about individual genomic sites into tensors. The feature mapper encapsulates choices about what features are potentially informative and how they should be converted to a fixed length vector. Mapped features are used to train a neural network (funnel architecture shown, see methods) that consists of a certain number (typically 1 to 6) of densely connected layers, with decreasing number of neurons. We stack a softmax output on top of the last layer of the funnel, where each softmax element represents the presence or absence of a specific combination of genotypes at the given genomic site. [Considering a diploid organism and two additional possible genotypes, the softmax has 17 elements (2^2+2^ + 1)]. The neural network model is trained by minimizing the difference between the model output and true labels. Hyper-parameters *r* and *c* are defined in methods.

### Training Protocols

In order to establish if semi-supervised methods are beneficial for this type of models and datasets, we defined four training protocols, two supervised and two semi-supervised:

- Supervised training (**Direct Supervised**). In this straightforward and popular training protocol, input features and labels over a training set are used to optimize the parameters of the model by minimizing a loss between model output and true labels.
- Supervised mixup training (**Mixup Supervised**). This approach was proposed to improve training of images [Zhang et al., 2017]. Supervised mixup trains the model using mixture examples constructed by combining two training examples in a linear fashion. This approach seems particularly relevant to training genotype callers since alignment data is the result of measuring a sample from a mixture of DNA molecules containing different genotypes (mixture of two alleles for most sites in a diploid genome). In this setting, mixup can be seen as a data augmentation strategy that constructs novel training examples using a linear combination of existing examples.
- Semi-supervised training with an adversarial autoencoder (**Semi-supervised AAE**). AAEs are a semi-supervised approach which has been shown to improve performance of models trained with small number of examples [Makhzani et al., 2015]. A semi-supervised training approach uses unlabeled examples in addition to labeled training examples. Here, we evaluated whether a semi-supervised AAE can improve genotype calling performance by taking advantage of unlabeled data from other alignments. This question differs from semi-supervised in small data regimes because we try to improve results when millions of training examples are already available. In this setting, it is unclear if adding unlabeled examples can still be of benefit.
- Semi-supervised mixup training (**Semi-supervised Mixup**). While mixup was initially described as a fully supervised approach, it is possible to adapt this approach in the semi-supervised setting by dreaming up labels. Here, we dream up labels for unlabeled examples by sampling from their distribution. We have evaluated two different sampling methods: either sampling from a uniform distribution, or sampling from the distribution of the labels observed in the training set.

### Hyper-parameter Optimization

Training a model using neural networks requires choosing a number of hyper-parameters. The performance of neural network models can critically depend on the choice of hyper-parameter, to the extent that some authors have shown that performance advantages obtained with new models can be achieved with baseline models when a similar hyper-parameter search is performed. Given the importance of hyper-parameter tuning, we report all results after tuning each method in a similar manner.

A random hyper-parameter search consists in choosing hyper-parameters randomly given a suitable range and distribution for each hyper-parameter. We performed these searches in two phases.

### Performance of semi-supervised approaches with 10,000 training examples

First, we generated 160 different random combinations of parameters and trained models with the first 10,000 training examples of the training sets. This produced the range of model performance shown in Figure 2. These results are surprising because semi-supervised methods are expected to perform best when few training examples are available. Instead, we observe that the two supervised methods perform best in these searches.

**Figure 2.**
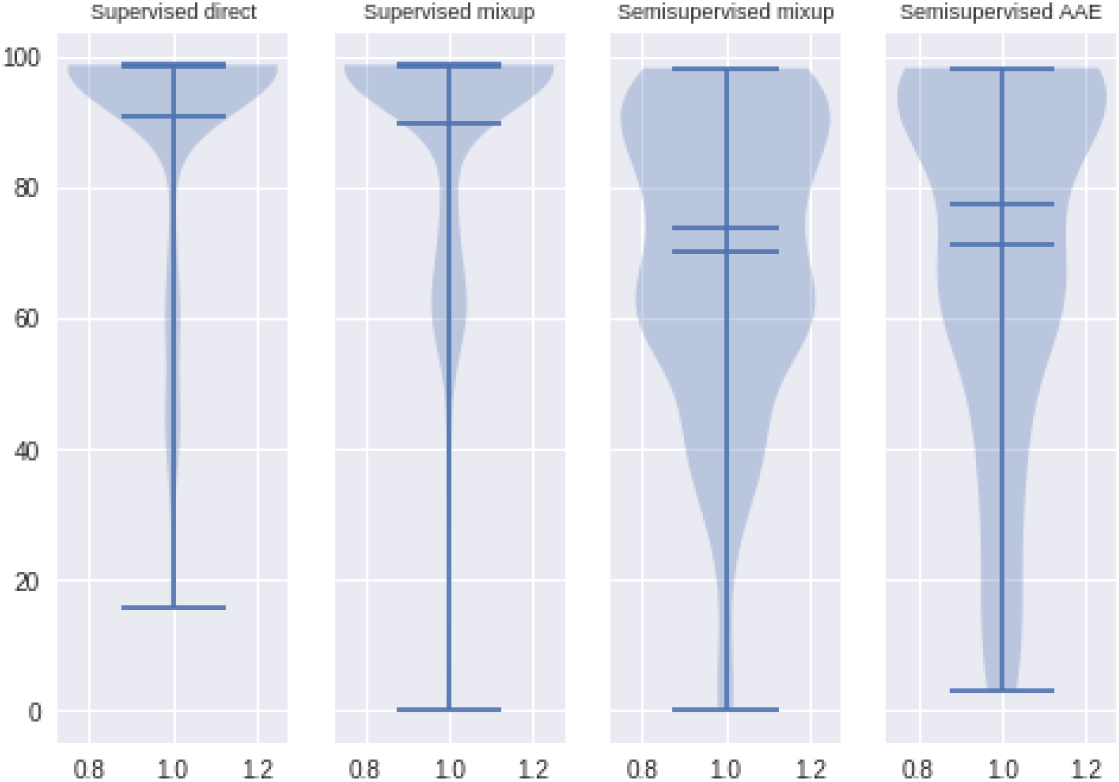
Validation accuracy for models produced in first phase of hyper-parameter search. We trained 160 models per training protocol, using the first 10,000 training examples. The plots show the accuracy of the model measured on the validation set.

### Domain Adaptation

The key assumption of model generalization is that the training dataset and the dataset where the model is used belong to the same data distribution. One possibility for the loss in performance observed when transferring a model trained on HG001 to HG002 is that the two datasets have different data distributions. Various domain adaptation techniques have been developed to mitigate the drop of performance in such situations. We evaluated the ability of the Adversarial Discriminative Domain Adaptation (ADDA) method to adjust the distribution of the data. Briefly, ADDA fine tunes the feature part of a model to make the features similarly distributed between the training set and the unlabeled set. The classifier part of the model is then used as is for classification. When we used ADDA to adapt a model to the HG002 data distribution, we found that the adapted model performed worse than the non-adapted model (see Table 1).

**Table 1.**
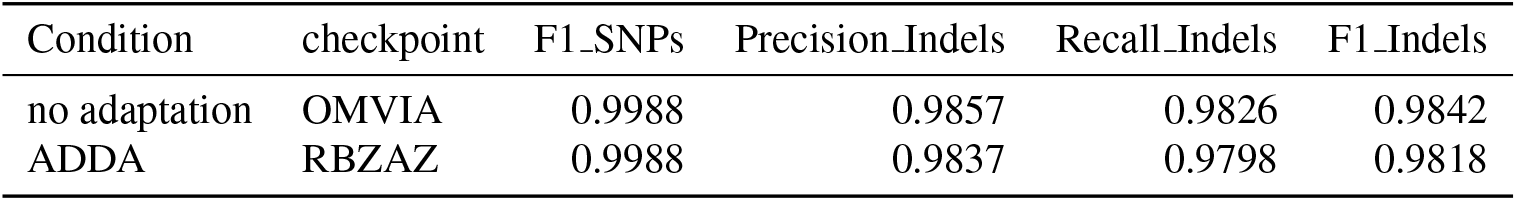
Domain Adaptation Results. These performance results are obtained in the confidence regions of the entire HG002 genome. The results indicate that the domain adaptation method tested is unable to improve performance for indels an independent sample, suggesting that systematic biases in data distributions are not responsible for the observed drop in performance.

### Performance with 3.9 million training examples

In a second phase of optimization, we use the entire training set, and choose random hyper-parameters from refined ranges. To refine each hyper-parameter range, we select the top three models for each training protocol and reduce the range to that observed in the top models. This process is illustrated in an online notebook [Onl]. We use these refined ranges to choose 16 sets of random hyper-parameters per training protocol. Figure 3 shows the accuracy (panel A) and indel F1 performance (panel B) of models trained with these optimized parameters. Figure 3 does not show the performance of the AAE training protocol because these models had non-competitive performance when trained on the entire training set (i.e., all accuracies were below 80%).

**Figure 3.**
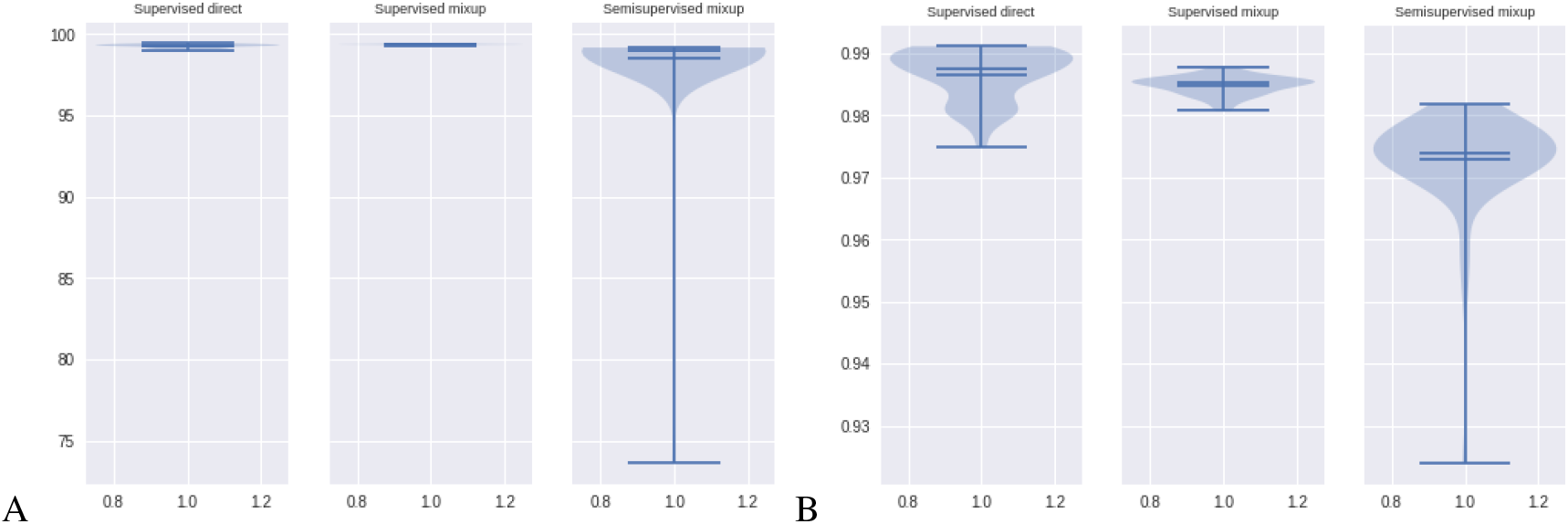
Validation accuracy for models produced in second phase of hyper-parameter search. We trained 64 models per training protocol, using the entire training set and optimized hyper-parameters. A) The plots show the accuracy of the models measured on the validation set. B) Violin plots showing indel F1 performance estimated on the validation set.

Table 2 presents the performance of the top 3 models, for each training protocol, measured on the validation set. We chose the best model by validation performance and evaluated its performance on the HG001 test set (test set performance shown in Table 3). Since the validation and test sets are fully independent, constructed from distinct sets of chromosomes of HG001, these data confirm that the models trained with GenotypeTensors generalize to new sequence variations and genomic contexts.

**Table 2.**
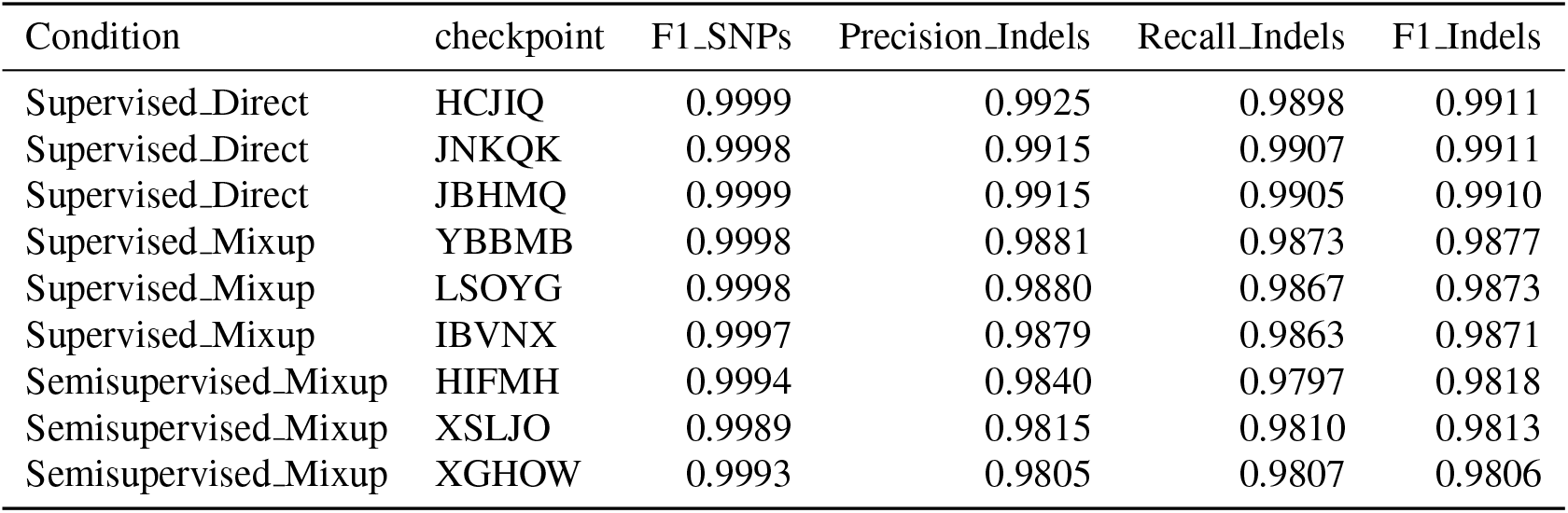
Top 3 HG001 validation performance for each training protocol.

| Condition | checkpoint | F1_SNPs | Precision_Indels | Recall_Indels | F1_Indels |
| --- | --- | --- | --- | --- | --- |
| Supervised_Direct | HCJIQ | 0.9999 | 0.9925 | 0.9898 | 0.9911 |
| Supervised_Direct | JNKQK | 0.9998 | 0.9915 | 0.9907 | 0.9911 |
| Supervised_Direct | JBHMQ | 0.9999 | 0.9915 | 0.9905 | 0.9910 |
| Supervised_Mixup | YBBMB | 0.9998 | 0.9881 | 0.9873 | 0.9877 |
| Supervised_Mixup | LSOYG | 0.9998 | 0.9880 | 0.9867 | 0.9873 |
| Supervised_Mixup | IBVNX | 0.9997 | 0.9879 | 0.9863 | 0.9871 |
| Semisupervised_Mixup | HIFMH | 0.9994 | 0.9840 | 0.9797 | 0.9818 |
| Semisupervised_Mixup | XSLJO | 0.9989 | 0.9815 | 0.9810 | 0.9813 |
| Semisupervised_Mixup | XGHOW | 0.9993 | 0.9805 | 0.9807 | 0.9806 |

**Table 3.** Best performance on HG001 test set across conditions. The top-model selected using validation performance (see Table 2) is evaluated on the test set composed of different chromosomes than the training and validation set.

| Condition | checkpoint | F1_SNPs | Precision_Indels | Recall_Indels | F1_Indels |
| --- | --- | --- | --- | --- | --- |
| Supervised_Direct | HCJIQ | 0.9997 | 0.9939 | 0.9936 | 0.9938 |
| Supervised_Mixup | YBBMB | 0.9997 | 0.9901 | 0.9896 | 0.9899 |
| Semisupervised_Mixup | HIFMH | 0.9995 | 0.9860 | 0.9849 | 0.9855 |

In order to test how well the models generalize to new genomes, and to compare the performance of our approach to other studies, we tested these models with the HG002 sample. Table 4 presents these results and shows that our approach remains strongly predictive on HG002, and substantially outperforms Clairvoyante [Luo et al., 2018]. We also compare the performance of three models trained with the direct supervised protocol when measured on the validation and test sets of HG001 and HG002. These data indicate that the model generalizes well (Table 5), but suggests that neither the validation or test sets are fully representative of the complexity of indels and genomic context over an entire genome (compare performance in test set in HG002 to performance on entire genome HG002, Table 4).

**Table 4.** Performance of top 3 performing models trained on HG001 and tested on HG002 (entire genome). The table compares the performance of models trained with our approach (best 3 of 64 hyperparameter search models) to the performance of models trained with Clairvoyante [Luo et al., 2018]. Since the Clairvoyante preprint only reported F1 estimated across both SNPs and indels, we report F1_Full for direct comparison. All of the top 3 models trained with our approach outperform the Clairvoyante model when tested on the same dataset. While the difference in performance may appear small when considering the F1_Full metric, these small changes reflect large indel F1 performance differences. Among the top 8 models for each of our approach, the one with the closest performance to 0.9952 obtained an F1 indel of 0.9685, while the top performing model reached 0.9842.

| Condition | checkpoint | F1_Full | F1_SNPs | Precision_Indels | Recall_Indels | F1_Indels |
| --- | --- | --- | --- | --- | --- | --- |
| Supervised_Direct | OMVIA | 0.9975 | 0.9988 | 0.9857 | 0.9826 | 0.9842 |
| Supervised_Direct | OVMVZ | 0.9974 | 0.9988 | 0.9858 | 0.9818 | 0.9838 |
| Supervised_Direct | QIHQJ | 0.9974 | 0.9989 | 0.9838 | 0.9823 | 0.9831 |
| Supervised_Mixup | MJOGP | 0.9967 | 0.9988 | 0.9779 | 0.9759 | 0.9769 |
| Supervised_Mixup | AWSTO | 0.9966 | 0.9987 | 0.9780 | 0.9750 | 0.9765 |
| Supervised_Mixup | ZVHAD | 0.9959 | 0.9982 | 0.9770 | 0.9722 | 0.9746 |
| Semisupervised_Mixup | CIKOE | 0.9960 | 0.9985 | 0.9708 | 0.9770 | 0.9739 |
| Semisupervised_Mixup | OTPKK | 0.9959 | 0.9984 | 0.9728 | 0.9723 | 0.9726 |
| Semisupervised_Mixup | PUYDA | 0.9959 | 0.9985 | 0.9723 | 0.9717 | 0.9720 |
| Clairvoyante (Luo et al.) | model1 | 0.9950 | NP | NP | NP | NP |
| Clairvoyante (Luo et al.) | model2 | 0.9952 | NP | NP | NP | NP |
| Clairvoyante (Luo et al.) | model3 | 0.9949 | NP | NP | NP | NP |

**Table 5.** Consistency of performance across validation and test sets HG001 vs HG002. The table compares the indel F1 performance of 3 models trained with supervised direct on the HG001 training set. Performance is shown for the validation and test sets of HG001 and HG002. Consistency of the predictions across test sets indicates that the models generalize well. A comparison of these indel performances with those reported in Table 4 suggests that the test set and validation set are not fully representative of the sites in an entire genome since test and validation performance over-estimates performance measured on the entire genome.

| Condition | checkpoint | HG001 val | HG001 test | HG002 val | HG002 test |
| --- | --- | --- | --- | --- | --- |
| Supervised_Direct | KTVNR | 0.9948 | 0.9943 | 0.9959 | 0.9956 |
| Supervised_Direct | LTPEC | 0.9948 | 0.9940 | 0.9959 | 0.9953 |
| Supervised_Direct | MWCTC | 0.9949 | 0.9934 | 0.9963 | 0.9951 |

To investigate this effect more carefully, we plot a precision recall curve for all predictions on the HG002 genome (Figure 4). This plot indicates that while most of the precision/recall curve behaves as expected for a strong model, some sites are erroneously predicted with the wrong genotype with high probability (region of the curve marked with an arrow). We found that this result is representative of the neural network models that we have trained, which when tested on sites with characteristics that they may not have encountered in the training set can fail with high-probability. Inspection of some of these errors suggest that most of them are confusion among homozygote and heterozygote calls. While we cannot rule out that these errors are due to issues remaining in our software implementation, these results suggest that strong indel models may require training with several genomes in the training set in order to capture enough indel diversity that such errors on new samples are minimized.

**Figure 4.**
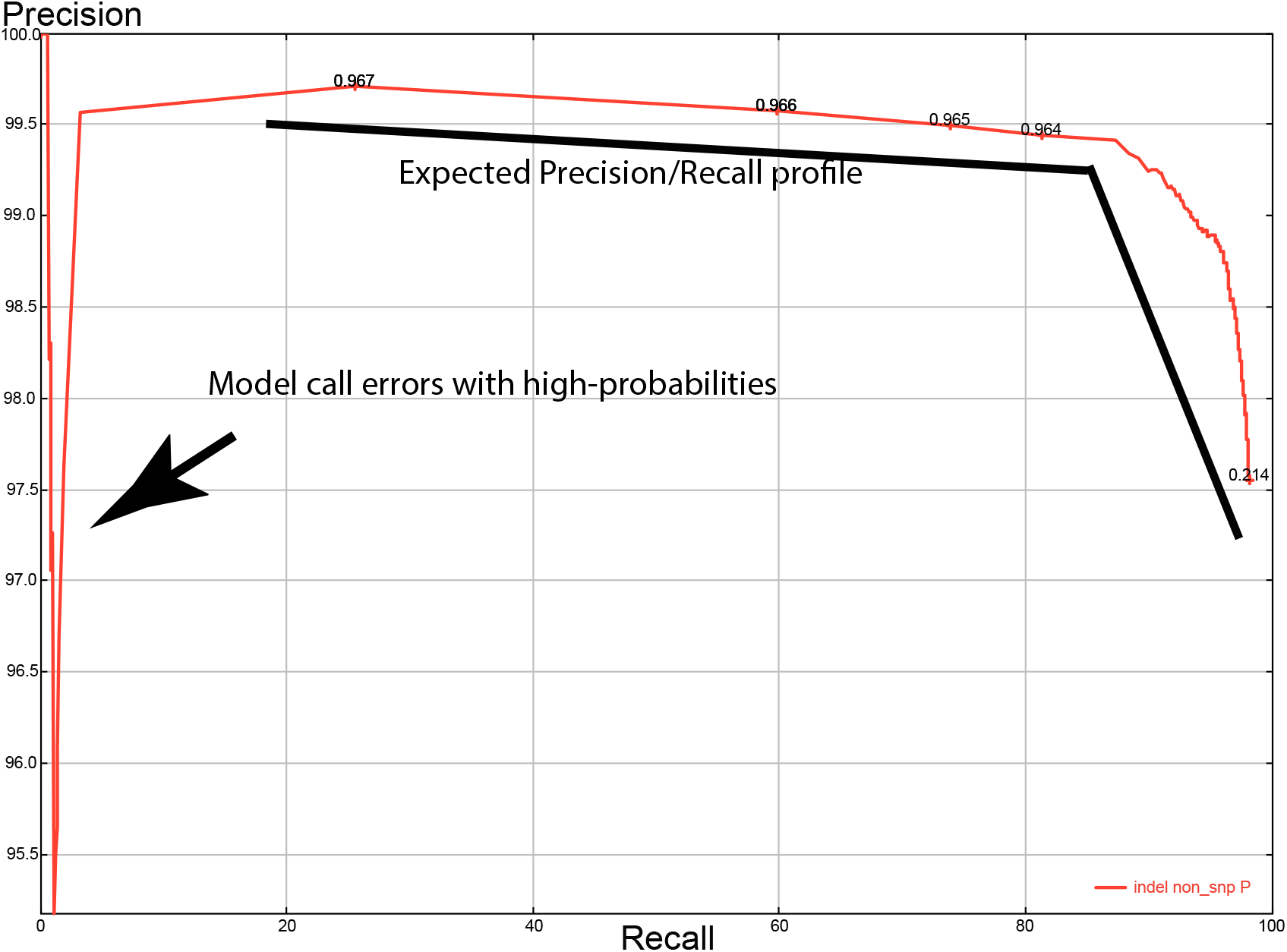
Precision Recall Curve for entire HG002 genome. This figure shows the precision recall curve for a model that performs well on the HG002 test set, but fails on some indel sites of HG002 with high-probability. Most of the curve is well behaved, but some sites yield indel genotype errors with very high probability (see arrow). This result is not an outlier, but instead typical of failure modes we have repeatedly observed when evaluating neural networks for genotype calling. Upon inspection, most of the errors are confusion between heterozygote or homozygote calls (i.e., one of the alleles is called correctly, but the other not, resulting in an overall incorrect call).

### Impact of training with sites in confidence regions only

Our preferred training protocol is to train with all sites of the genome, in order to avoid biasing results for possibly easier sites that may be contained in the GIAB confidence regions of the genome. Here, we tested this assumption by training models exclusively with sites contained in the GIAB confidence regions. We performed a random hyper-parameter search on HG001 and tested the top model on HG002. Table 6 presents the performance obtained on HG002 and shows that while validation performance is very high (accuracy reaching 99.9681 on the validation set on HG001), the model (MOMWU) performs worse on HG002 than when models are trained with sites both inside and outside of confidence regions (top model shown for reference: OMVIA).

**Table 6.**
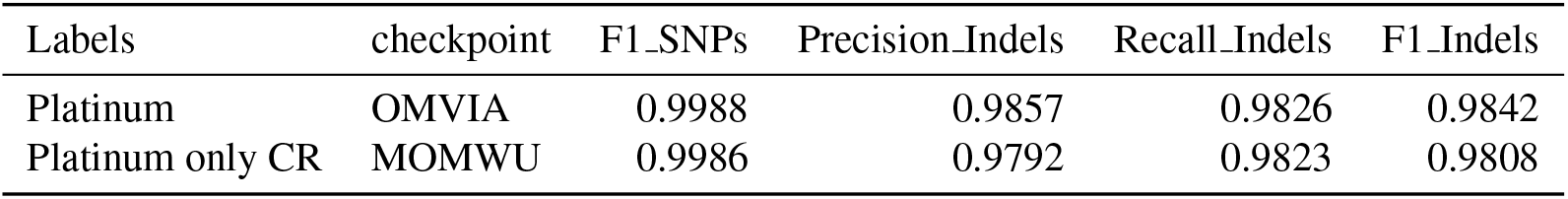
Training only with confidence regions (CR). We train 64 models, random hyper-parameter search using only sites contained in GIAB confidence regions. The validation performance of the top model measured on HG001 is shown (checkpoint: MOMWU).

| Labels | checkpoint | F1_SNPs | Precision_Indels | Recall_Indels | F1_Indels |
| --- | --- | --- | --- | --- | --- |
| Platinum | OMVIA | 0.9988 | 0.9857 | 0.9826 | 0.9842 |
| Platinum only CR | MOMWU | 0.9986 | 0.9792 | 0.9823 | 0.9808 |

### Impact of GIAB genotype call improvements

Models trained until this point were trained with genotype calls from the platinum genome [Eberle et al., 2016]. Since genotype calls have been refined over the years by the GIAB consortium, we trained a model with version 3.3.2 of the GIAB calls to determine the impact of true label refinements on the performance of our models. Table 7 presents these results and shows that small performance improvements can be obtained by using more recent versions of the true labels, but that our training protocol is quite robust to some errors in genotypes (i.e., the platinum calls contain more errors than the GIAB 3.3.2 calls).

**Table 7.** Impact of genotype label improvements. We compare the performance (HG002 entire genome) of the top model obtained when training with platinum labels or with the latest release of the GIAB genotypes for HG001 (best of 64 models trained with random hyper-parameter search shown for each condition). The comparison shows that a small performance improvement is obtained by using more recent versions of the genotype calls, but also clearly confirms that our approach can learn despite errors typically found in the genotype calls used in the training set.

| Labels | checkpoint | F1_SNPs | Precision_Indels | Recall_Indels | F1_Indels |
| --- | --- | --- | --- | --- | --- |
| Platinum | OMVIA | 0.9988 | 0.9857 | 0.9826 | 0.9842 |
| GIAB 3.3.2 | KTVNR | 0.9989 | 0.9865 | 0.9840 | 0.9852 |

### Computational Performance

The approaches we presented are implemented in the GenotypeTensors project, developed as an open-source project available at https://github.com/CampagneLaboratory/GenotypeTensors. The project leverages the popular PyTorch deep-learning framework. Various optimizations are implemented in GenotypeTensors to train models efficiently. Table 8 presents the time to train one epoch for HG001 (with 3,923,305 training examples) with GenotypeTensors. Timing is dependent on the numbers of parallel processes used to load data from disk and feed it to the GPU, with more workers resulting in faster training. The results shown in table 8 were obtained when caching data on a solid state drive.

**Table 8.** Computational Performance. The table shows the number of seconds needed to train a top-performing model for a single epoch using a representative HG001 dataset. The time required to train an epoch (one pass through the entire dataset) is determined by the time it takes to load the data from disk and transfer it to the GPU. Loading data in parallel is supported by PyTorch and adding more workers results in faster training. Performance gains plateaus at about 20-30 workers.

| Number of parallel workers | Time to train an epoch (secs) |
| --- | --- |
| 6 | 204 |
| 10 | 136 |
| 20 | 63 |
| 30 | 53 |

Another practical aspect of computational performance is the ability to quickly determine optimal hyper-parameters for training a model. As illustrated in Figure 3 (Panel B), the performance of models varies substantially with the choice of hyper-parameters, so it is important to determine a set of optimal hyper-parameters for a dataset. Since the data loading time is the bottleneck when training models, we have developed code to load data once and train several models initialized with different hyper-parameters (see Methods for details). This approach makes it possible to train up to 64 models in parallel on the same GPU, and greatly reduces the computational burden of performing random hyper-parameter searches. With this approach, we are able to train 64 models for about 10 epochs on a single GPU in about 6-8 hours (without this approach, training 16 models on 8 GPUs independently can be done in a few days). We do not provide specific timing for this process because the search time depends on the range of hyper-parameters explored (i.e., the number of parameters optimized for each model included in a search will influence the runtime). However, this approach makes hyper-parameter selection much more practical than training models independently.

### Cloud applications

We have developed DNANexus cloud applications to help process alignments and generate training and evaluation datasets, or to call genotypes across whole genomes with trained models. The source code of these applications is available at https://github.com/CampagneLaboratory/variationanalysis-apps.

## DISCUSSION

We have presented an efficient approach to train neural network methods for calling genotypes. The method is orders of magnitude more efficient than DeepVariant because it converts genomic sites to a few thousand features and uses a feed forward network instead of a convolutional architecture trained from images. While our approach is computationally efficient, it has not yielded models competitive with the performance reported for DeepVariant on HG002, or other more traditional approaches evaluated in the PrecisionFDA challenge. For instance, the best performance reported on the entire genome of HG002 in the blind evaluation of PrecisionFDA (Truth challenge) reached an indel F1 of 0.9904 (99.4%). This compares to our best model reaching an indel F1 of 0.9852. In the same challenge, DeepVariant reached an indel F1 of 0.9898. While DeepVariant has since improved performance on HG002, the performance statistics are no longer directly comparable to our results because recent DeepVariant models are trained with several genomes, in contrast to the HG001 training protocol we use for evaluation in this study.

Given the loss of performance outside of the test and validation sets that we observed in Figure 4, it is likely that adding more genomes to the training set in order to optimize HG002 performance would yield biased performance estimates (i.e., stronger performance would be observed on HG002 than on a randomly selected genome, where distinct high probability errors similar to those shown in Figure 4 could manifest).

Given these results, it is unclear if the superior performance of DeepVariant is due to the convolutional architecture used in the DeepVariant models, which gives the model access to data about the context of each specific genomic site (e.g., DeepVariant provides data about 150bp around each site) and represents a tradeoff between computational efficiency and model performance, or if the implementation of our approach could be further improved to match deep-variant like model performance at a fraction of the computational cost.

A new tool, called Clairvoyante, was recently evaluated in a preprint [Luo et al., 2018]. This tool implements a neural-network based method for calling SNPs and indels in short and long-read data. Clairvoyante is described as a convolutional network and therefore also must convert alignment data to three dimensional tensors. The preprint provides data showing that Clairvoyante is faster than DeepVariant. Interestingly, no comparison, either computational efficiency or call performance, is provided with our earlier project, variation analysis (whose performance kept improving since the release of our initial preprint [Torracinta and Campagne, 2016]). We have included Clairvoyante in our evaluation in this manuscript and shown that all the training protocols we tested outperform Clairvoyante on HG002 by a wide margin when considering indel performance (see Table 4). Our approach has an advantage as well for computational performance since (1) we can train an epoch with HG001 in less than a minute (Table 8) when Clairvoyante needs 170 seconds [Luo et al., 2018] and (2) we can perform a hyper-parameter search with 64 models in the time it takes to train one Clairvoyant model. Regarding indel performance, as we have noted in the caption of Table 4, [Luo et al., 2018] presented F1 performance evaluated across both SNPs and indels. This made comparison with prior studies that separate SNP and indel performance difficult. We find this presentation confusing because it could mislead readers who quickly glance at numerical performance metrics without realizing that Full_F1 (Indel+SNP) can appear numerically competitive with indel F1, when it is far from competitive. The comparison we provide in Table 4 should clarify how the performance of Clairvoyante compares to prior neural network methods. Additionally, we note that there is no reason that our approach, as described previously and in this manuscript, would be expected to be inferior to the approach implemented in Clairvoyante for long-read datasets. Indeed long-read technology does not fundamentally change what type of features are informative when considering a specific genomic site after the reads have been aligned to the genome. We therefore expect that simply performing a hyper-parameter search with long-read data with the open-source software that we provide will yield strong performance both for long-read and short-read sequencing datasets.

Clairvoyante uses a convolutional architecture similar to DeepVariant, but fails to match the performance of either our simpler single-site models, or DeepVariant models. Therefore, it seems unlikely that a convolutional architecture is sufficient to explain the difference in performance observed between our approaches and DeepVariant.

We hypothesize that so-far unnoticed software implementation problems in available code bases, and/or insufficient hyper-parameter tuning for Clairvoyante, could be responsible for the differences observed in model performance, rather than being driven only by differences in model architecture and differences in representation of aligned reads in each genomic context.

We have tested a domain adaptation approach and two semi-supervised approaches for genotype calling. In both cases, we found that these approaches, which have been successful in other domains, failed to improve the generalization of genotype caller models for indels or SNPs. This point supports the notion that data distribution differences are not primarily responsible for variations in performance from one individual sample to another.

## METHODS

### Datasets

HG001 and HG002 read files were obtained from PrecisionFDA (https://precision.fda.gov/).

### Reads Processing

Read files were uploaded to DNANexus and processed as follows. Alignments to the genome were done with BWA (DNANexus application) with default parameters. Alignments were further processed with DNANexus applications developed by our laboratory: alignments were processed with HaplotypeCaller (GATK3) to realign SNPs around indels and reduce the size of the training set to regions likely to contain non-reference calls. Realigned BAM files were converted to Goby alignment format, then to Sequence Base Information (SBI) format. Finally, SBI files were mapped to tensors with the export-tensors tool distributed in the variationanalysis project. Tensors files were then downloaded to the laboratory local server to train models.

### True Genotype Labels

Unless otherwise noted under Results, we used the platinum genome VCF file version 2017-1.0 as training data for NA12878. SNP and indel performance were measured against annotations from the GIAB consortium by restricting evaluation to confidence region.

### Training Neural Network Models

Models were developed with the PyTorch framework (http://pytorch.org/), version 0.3.1. PyTorch was selected because it implements automatic differentiation and is a popular framework with strong community support. All models were trained with a batch size of 1024 examples. The order of training examples is shuffled at each epoch. We trained models on a Linux server with 8 GTX 1080Ti GPUs (11GB of RAM per GPU, 256GB of RAM for the server), but we note that single model training can be conducted on a single GPU overnight in a couple of hours. Hyper-parameter search for 64 models can be conducted on a single GPU overnight (10 epochs).

### Hyper-parameter searches

We implemented a fast random hyper-parameter search approach in GenotypeTensors. Since our models occupy only a few megabytes of GPU memory, we are able to train 64 different models (each initialized with different hyper-parameters) at the same time on a single GPU. Briefly, we load tensors for a mini-batch and then perform forward, backward and parameter updates for a single model in independent threads. Because most of the computationally expensive work is done on the GPU, the python Global Interpreter Lock is not a major bottleneck and tens of models can be trained in the same time that a single model would be trained if done sequentially. When all models are trained with a mini-batch, another batch of tensors is loaded and training continues until all batches have been used. Testing can similarly be done for 64 models in different threads. An important advantage of this approach is that a batch of data is loaded only once. This provides an important speedup when the training set is large and disk or solid state drive random access is needed to load batches. This approach is implemented in https://github.com/CampagneLaboratory/GenotypeTensors/blob/master/src/org/campagnelab/dl/genotypetensors/autoencoder/SearchHyperparameters.py.

### Best model hyperparameters

This section provides the command lines showing hyper-parameters used to train a few of the top-performing models discussed under Results. Note that using these command lines to train a model requires to add the path to the dataset to use for training/validation. This is done by adding an argument of the form –problem genotyping:NA12878-realigned-2018-01-30, where NA12878-realigned-2018-01-30 is the path to a dataset. Adding the option –num-workers 6 on a machine with 6 cores will also result in faster training by loading data in parallel.

~~~
OMVIA:
train-autoencoder.sh --mode supervised_funnel_genotypes \
--L2 9.033815206858845E-5 --dropout-probability 0 \
--optimizer adagrad --lr 0.009161936 \
--reduction-rate 0.39535806 \
--model-capacity 1.0 --num-layers 2 \
--indel-weight-factor 2.0382204 \
--max-epochs 100 --epoch-min-accuracy 80.0 \
--mini-batch-size 1024 --epsilon-label-smoothing 0.0958678 \
--seed 3453641268241957507
~~~

~~~
KTVNR:
train-autoencoder.sh --mode supervised_funnel_genotypes \
--L2 9.125417163009775E-5 \
--dropout-probability 0 --optimizer adagrad \
--lr 0.006005622 --reduction-rate 0.4112884 \
--model-capacity 1.0 --num-layers 2 \
--indel-weight-factor 9.482289 \
--max-epochs 100 --epoch-min-accuracy 80.0 \
--mini-batch-size 1024 --epsilon-label-smoothing 0.030408645 \
  --seed 950779171533190880
~~~

#### Funnel Architecture (FA)

Funnel models were formulated as *n* fully connected layers with BatchNormalization and with RELU/SELU activation and a fully connected output layers with soft-max activation. The number of output layers *n* is a hyper-parameter of the architecture that we optimized (see hyper-parameter search). The first dense inner layer contains *c* times the number of input features (we call *c* the model capacity). The number of neurons of each layer is calculated by applying a reduction factor *r* to the number of input to the layer. When *r* < 1, the model looks like a funnel and progressively reduces the number of neurons until the softmax layer. The exact model architecture used is encoded in the class called `org.campagnelab.dl.genotypetensors.autoencoder.genotype_softmax_classifier.GenotypeSoftmaxClassifer` distributed in the Geno-typeTensors project.

#### Adversarial AutoEncoder Architecture (AAEA)

Adversarial AutoEncoder is implemented as described in [Makhzani et al., 2015]. This architecture is composed of an encoder, made of *n* fully connected layers of decreasing number of neurons. The encoder reduces the number of features progressively to a configurable number *z*. A decoder implements the reverse transformation to reconstitute the input features from the encoded representation. A discriminator loss is used to make each of the *z* features approximately normally distributed (zero mean and unit standard deviation). The decoder and encoder are trained to reconstruct unlabeled examples while semi-supervision is obtained by training the encoder to produce *z* features that can predict the labels of training set examples.

#### Direct Supervised Training

Direct supervised training optimizes the supervised loss using an Adagrad optimizer with an L2 loss (an optimized hyper-parameter). We use the PyTorch MultiLabelSoftMarginLoss.

#### Mixup Supervised Training

Mixup supervised training optimizes the supervised loss using an Adagrad optimizer with an L2 loss (an optimized hyper-parameter). In contrast to direct supervised training, mixup training uses two training examples and combines their features and labels in a linear manner. *x_mixup_* = *α.x*_1_ + (1 − *α*).*x*_2_, *y_mixup_* = *α.y*_1_ + (1 − *α*).*y*_2_ We use the PyTorch MultiLabelSoftMarginLoss.

#### Mixup Semisupervised Training

This training procedure is similar to mixup supervised training, but instead of using two training examples, we use one training example and one unlabeled example. The label of the unlabeled example is sampled from a distribution. Two sampling approaches are implemented: uniform (each genotype class is equiprobable), sampling (each genotype class is sampled with probability the frequency of the genotype in the training set).

#### Label smoothing

During all types of training, we use label smoothing with configurable ε [Goodfellow et al., 2016] (section 7.5.1). An *ε* of 0 means that labels are not smoothed. Label smoothing is also consistently applied in semi-supervised training schemes when labels are sampled from a distribution.

### Performance Metrics

Accuracy is measured as TP/(TP+FP) on the softmax prediction for each site using the GenotypeTensors software. SNP and indel F-1 are estimated with the Real Time Genomics software tool rtg vcfeval, from genotypes calls made with variationanalysis and GenotypeTensors. SNP and indel performance is estimated independently, using baselines restricted to each type of variation. The following command line is used for rtg to estimate performance for SNPs:

~~~
  rtg vcfeval --baseline=GOLD-confident-chr-snps.vcf.gz \
      -c direct-best-WHVBC-CNG-NA12878-realigned-2018-01-30-test-\
genotypes-sorted.vcf-snps.vcf.gz \
      -o output-4092/snp --template=hs37d5_all_chr_CNG.sdf \
      --evaluation-regions=GIAB-NA12878-confident-regions-chr.bed.gz \
      --bed-regions=direct-best-WHVBC-CNG-NA12878-\
realigned-2018-01-30-test-observed-regions.bed-sorted.bed.gz \
      --vcf-score-field=P --sort-order=descending
~~~

## ACKNOWLEDGMENTS

We thank Rémi Torracinta for his contributions to an earlier version of the project.

